# TCF7L2 regulates postmitotic differentiation programs and excitability patterns in the thalamus

**DOI:** 10.1101/515874

**Authors:** Marcin Andrzej Lipiec, Kamil Koziński, Tomasz Zajkowski, Joanna Bem, Joanna Urban-Ciećko, Michał Dąbrowski, Chaitali Chakraborty, Łukasz Mateusz Szewczyk, Angel Toval, José Luis Ferran, Andrzej Nagalski, Marta Barbara Wiśniewska

**Author notes:** Correspondence should be addressed to Marta B. Wiśniewska. These authors contributed equally (Kamil Koziński, Tomasz Zajkowski, Joanna Bem).

## Abstract

Neuronal phenotypes are controlled by terminal selector transcription factors in invertebrates, but few examples of such regulators have been provided in vertebrates. TCF7L2 has been identified as a regulator of efferent outgrowth in the thalamus and habenula. We used a complete and conditional knockout of *Tcf7l2* in mice to investigate the hypothesis that TCF7L2 plays a dual role in thalamic neuron differentiation and functions as a terminal selector. Connectivity and cell clustering was disrupted in the thalamo-habenular region in *Tcf7l2^-/-^* embryos. The expression of subregional thalamic and habenular transcription factors was lost and region-specific cell migration and axon guidance genes were downregulated. In mice with postnatal *Tcf7l2* knockout, the induction of genes that confer terminal electrophysiological features of thalamic neurons was impaired. Many of these genes proved to be TCF7L2 direct targets. The role of TCF7L2 in thalamic terminal selection was functionally confirmed by impaired firing modes in thalamic neurons in the mutant mice. These data corroborate the existence of master regulators in the vertebrate brain that maintain regional transcriptional network, control stage-specific genetic programs and induce terminal selection.

**Statement:** The study describes a role of TCF7L2 in neuronal differentiation of thalamic glutamatergic neurons at two developmental stages, highlighting its involvement in the postnatal establishment of critical thalamic electrophysiological features.

## Introduction

Studies of invertebrates have shown that terminal differentiation gene batteries in individual classes of neurons are induced and maintained by specific transcription factors that are expressed throughout the life, called terminal selectors (Hobert and Kratsios, 2019). Regulatory strategies of postmitotic maturation and terminal selection in vertebrates are unclear because only a few terminal selectors have been identified to date (Cho et al., 2014; Flames and Hobert, 2009; Kadkhodaei et al., 2009; Liu et al., 2010; Lodato et al., 2014; Wyler et al., 2016). The thalamus and habenula are glutamatergic regions that derive from a single progenitor domain, prosomere 2 (Watson et al., 2012). The thalamus is a sensory relay centre, and part of the cortico-subcortical loops that process sensorimotor information and produce goal-directed behaviours (Sherman, 2017). The habenula controls reward- and aversion-driven behaviours by connecting cortical and subcortical regions with the monoamine system in the brainstem (Benekareddy et al., 2018; Hikosaka, 2010). During postmitotic differentiation, thalamic and habenular neurons segregate into discrete nuclei (Shi et al., 2017; Wong et al., 2018), develop variety of subregional identities (Nakagawa, 2019; Phillips et al., 2018), extend axons toward their targets (Hikosaka et al., 2008; López-Bendito, 2018), and acquire electrophysiological characteristics postnatally (Yuge et al., 2011). Knowledge of mechanisms that control postmitotic development in this region is important, because its functional dysconnectivity, which possibly originates from the period of postmitotic maturation, is implicated in schizophrenia, autism and other mental disorders (Browne et al., 2018; Steullet, 2019; Whiting et al., 2018; Woodward et al., 2017).

The network of postmitotically induced transcription factors that regulates the maturation of prosomere 2 neurons has only begun to be deciphered. *Gbx2* and *Pou4f1* are early postmitotic markers of thalamic and habenular neurons, respectively. GBX2 plays a transient regulatory role in the initial acquisition of thalamic molecular identities and thalamocortical axon guidance, and then its expression is downregulated in the majority of the thalamus (Chatterjee et al., 2012; Chen et al., 2009; Li et al., 2012; Mallika et al., 2015; Miyashita-Lin et al., 1999). In contrast, POU4F1 (alias BRN3A) is not essential for the growth of habenular axons, but maintains the expression of the glutamate transporter gene *Vglut1/Slc17a7* and other habenula-abundant genes in adult, though its impact on electrophysiological responses of habenular neurons was not tested (Quina et al., 2009; Serrano-Saiz et al., 2018). Subregional transcription factors RORA and FOXP2 regulate some aspects of postmitotic differentiation in thalamic subregions during embryogenesis (Ebisu et al., 2016; Quina et al., 2009; Vitalis et al., 2017), but their role in terminal differentiation in this region was not investigated.

*Tcf7l2*, a risk gene for schizophrenia and autism (Bem et al., 2019) that encodes a member of the LEF1/TCF transcription factor family (Cadigan and Waterman, 2012), is the only shared marker of prosomere 2 neurons (Nagalski et al., 2013; Nagalski et al., 2016). TCF7L2 expression is maintained throughout life and TCF7L2 motifs are overrepresented in putative enhancers of adult thalamus-enriched genes (Nagalski et al., 2016; Wisniewska et al., 2012), suggesting that this factor can be prosomere 2 terminal selector, but its function was tested only during early postmitotic period in zebrafish (Beretta et al., 2013; Husken et al., 2014) and mice (Lee et al., 2017). In *Tcf7l2^-/-^* mouse embryos, some markers were misexpressed in the region of prosomere 2, and the formation of axonal tracts was disrupted. The role of TCF7L2 in later developmental stages has not been investigated.

The present study used complete and conditional knockout mice to explore the role of *Tcf7l2* in postmitotic anatomical maturation, maintenance of molecular diversification, adoption of neurotransmitter identity, and postnatal acquisition of electrophysiological features in the thalamus and habenula. We show that TCF7L2 orchestrates the overall morphological differentiation process in this region by regulating stage-specific gene expression directly or via sub-regional transcription factors. We also report that TCF7L2 functions as a terminal selector of postnatally-induced thalamic electrophysiological characteristics but not glutamatergic (VGLUT2) identity.

## Results

### Generation of mice with the complete and conditional knockouts of Tcf7l2

*Tcf7l2* expression in prosomere 2 is induced in mice during neurogenesis (Cho and Dressler, 1998) and maintained in the thalamus and medial habenula throughout life (Nagalski et al., 2013). On E12.5, we observed high levels of TCF7L2 protein in the superficial portion of the thalamus and habenula, which is populated by postmitotic neurons (Fig. 1A). At late gestation, high TCF7L2 levels were observed in the entire caudal thalamus (hereinafter referred to as the thalamus; a glutamatergic domain) and medial habenula, and lower levels were in the rostral thalamus (rTh, a GABAergic domain) and lateral habenula (Fig. 1B).

**Figure 1.**
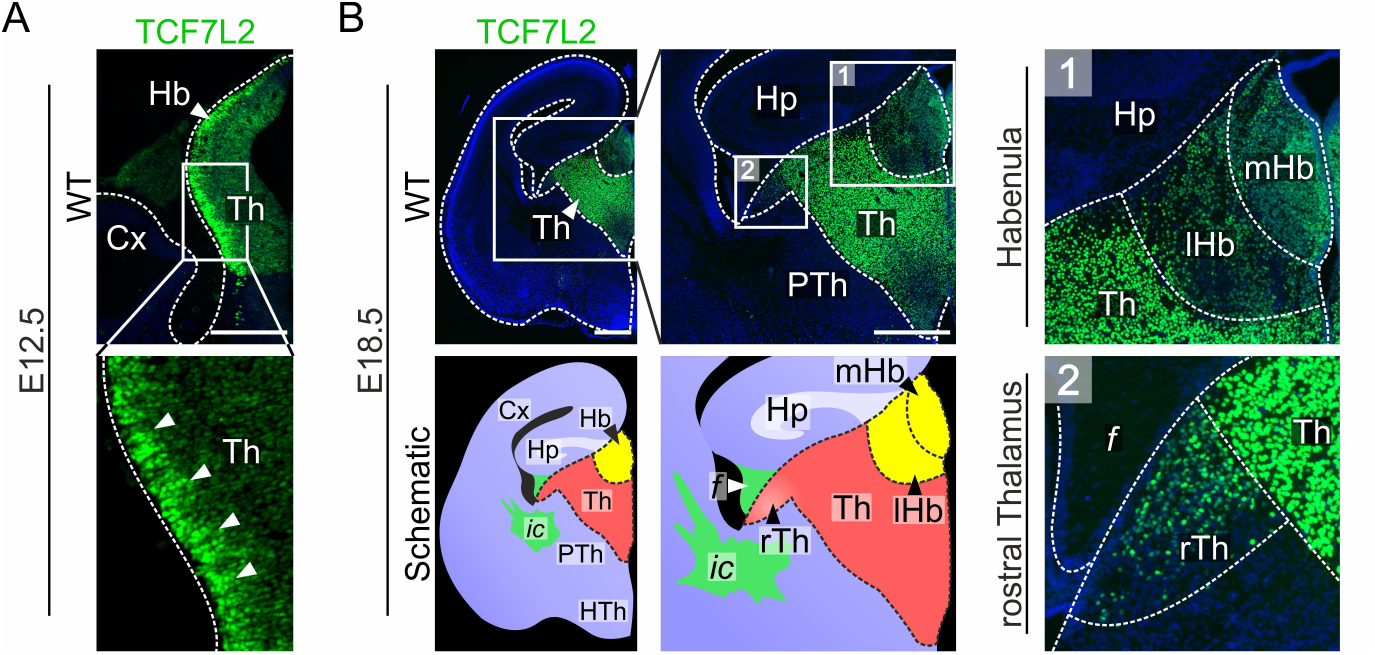
TCF7L2 protein levels are high in postmitotic neurons in the thalamo-habenular region. Immunofluorescent staining of TCF7L2 in E12.5 **(A)** and E18.5 **(B)** coronal brain sections. On E12.5, TCF7L2 is already detected in prosomere 2, but its level is high in the mantle zone (marked with white arrowheads on zoom-in). On E18.5, the thalamo-habenular region is marked by the abundance of TCF7L2. Local differences in TCF7L2 levels and cell densities enable to discriminate habenula **(1)** and rostral thalamus **(2)** from the central thalamic region. Cx, cortex; f, fornix; Hb, habenula; Hp, hippocampus; HTh, hypothalamus; ic, internal capsule; lHb, lateral habenula; mHb, medial habenula; PTh, prethalamus; rTh, rostral thalamus; Th, thalamus. Scale bars represent 0.5 mm.

To investigate the role of TCF7L2 as a selector of morphological as well electrophysiological characteristics of prosomere 2, we used two knockout models in mice. The complete knockout of *Tcf7l2* (*Tcf7l2*^tm1a/tm1a^, hereinafter referred to as *Tcf7l2*^-/-^) was generated by an insertion of the *tm1a(KOMP)Wtsi* allele with *lacZ* cassette upstream of the critical 6^th^ exon of the *Tcf7l2* gene (Fig. 2A). *Tcf7l2*^tm1a^ allele led to the lack of TCF7L2 protein, confirmed by immunostaining and Western blot (Fig. 2B-C), and ectopic expression of β-galactosidase from the *lacZ* locus. *Tcf7l2*^-/-^ mice die after birth. To create a thalamus-specific postnatal knockout of *Tcf7l2* we first crossed *Tcf7l2^+/^*^tm1a^ mice with transgenic animals that expressed flippase and then with mice that expressed CRE recombinase from the Cholecystokinin (*Cck)* gene promoter. *Cck* is upregulated postnatally and is expressed specifically in the thalamus and pallium (Allen Brain Atlas, 2011; Nishimura et al., 2015). In the resulting *Cck*^Cre^:*Tcf7l2*^fl/fl^ mice, TCF7L2 was absent in most thalamic nuclei in adult, but was maintained in the medial habenula (Fig. 2D-F). Cck-driven knockout of *Tcf7l2* was induced postnatally and completed in the thalamus by P14 (Fig. 2G-H).

**Figure 2.**
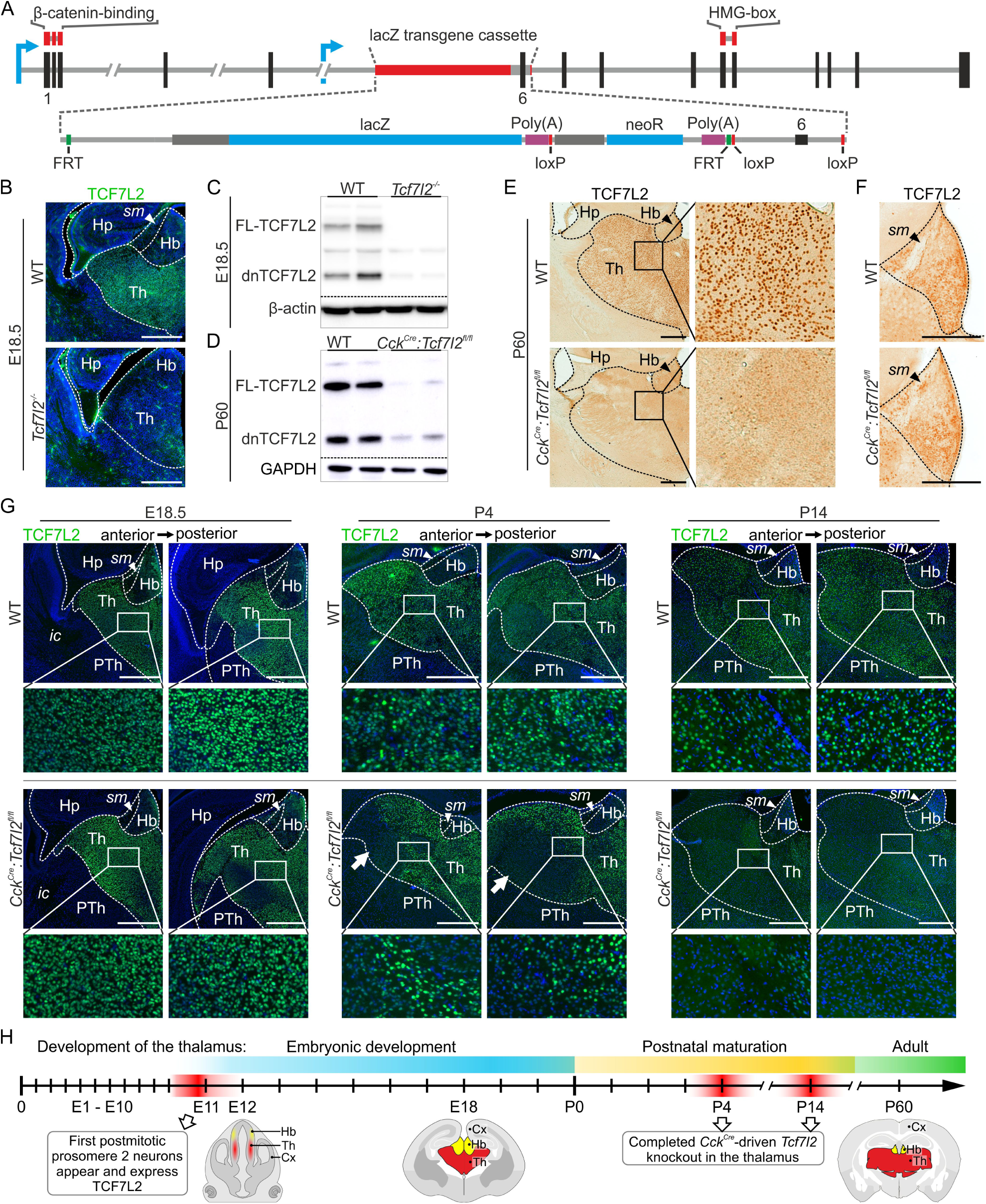
Generation of *Tcf7l2^-/-^* and *Cck^Cre^:Tcf7l2^fl/fl^* mouse strains. (A) Schematic representation of *Tcf7l2^tm1a^* allele generated by EUCOMM, in which a trap cassette with the *lacZ* and *neoR* elements was inserted upstream of the critical exon 6 of the *Tcf7l2* gene. Exons and introns are represented by vertical black and horizontal gray lines, respectively. Blue arrows indicate transcription start sites. Regions that encode the β-catenin binding domain and HMG-box are marked by red lines above the exons. **(B)** Immunofluorescent staining of TCF7L2 in coronal brain sections on E18.5 WT and *Tcf7l2^-/-^*. The expression of TCF7L2 protein is lost in *Tcf7l2^-/-^* embryos. **(C)** Western blot analysis of TCF7L2 protein expression in the thalamus in WT and *Tcf7l2^-/-^* mice on E18.5. Higher TCF7L2 bands correspond to the full-length protein (FL-TCF7L2), and the lower bands correspond to the truncated dominant negative isoform of TCF7L2 (dnTCF7L2). **(D)** Western blot analysis of TCF7L2 protein expression in the thalamus in WT (*Cck^Cre^:Tcf7l2^+/+^*) and *Cck^Cre^:Tcf7l2^fl/fl^* mice on P60. **(E)** DAB immunohistochemical staining of TCF7L2 in coronal brain sections on P60 WT and *Cck^Cre^:Tcf7l2^fl/fl^* mice. TCF7L2 is absent in most thalamic nuclei in adult *Cck^Cre^:Tcf7l2^fl/fl^* mice. **(F)** Zoomed-in view of the adult habenula, where TCF7L2 retains its presence in adult *Cck^Cre^:Tcf7l2^fl/fl^* mice. **(G)** Immunofluorescent staining of TCF7L2 in coronal brain sections in WT and *Cck^Cre^:Tcf7l2^fl/fl^* mice on E18.5, P4 and P14. Induction of the *Cck^Cre^*-driven *Tcf7l2* knockout occurs postnatally and by P4 the depletion of TCF7L2 protein is seen in the ventro-lateral regions of the thalamus (marked by white arrows). Progression of the *Cck^Cre^*-driven TCF7L2 depletion is completed in most thalamic nuclei by P14. **(H)** Simplified time course of the development of the thalamus with representative mouse brain schematics on E12.5; E18.5 and in adult. Cx, cortex; Hb, habenula; Hp, hippocampus; ic, internal capsule; PTh, prethalamus; sm, stria medullaris; Th, thalamus. Scale bars represent 0.5 mm.

### Normal neurogenesis but disrupted anatomy and connectivity of prosomere 2

To confirm that proliferation and neurogenesis occurred normally within prosomere 2 in *Tcf7l2*^-/-^ mice, we stained E12.5 brain sections with antibodies specific for the KI-67 antigen and TUJ1, markers of proliferating progenitors and young postmitotic neurons, respectively. Prosomere 2 was identified with *Tcf7l2* probe both in wild-type (WT) and knockout (KO) embryos, taking advantage of preserved expression from the targeted *Tcf7l2* locus (Fig. 3A). The complete knockout of *Tcf7l2* did not cause any apparent defects in proliferation or neurogenesis in prosomere 2 on E12.5 (Fig. 3B), consistently with previous results in another *Tcf7l2* knockout strain (Lee et al., 2017).

**Figure 3.**
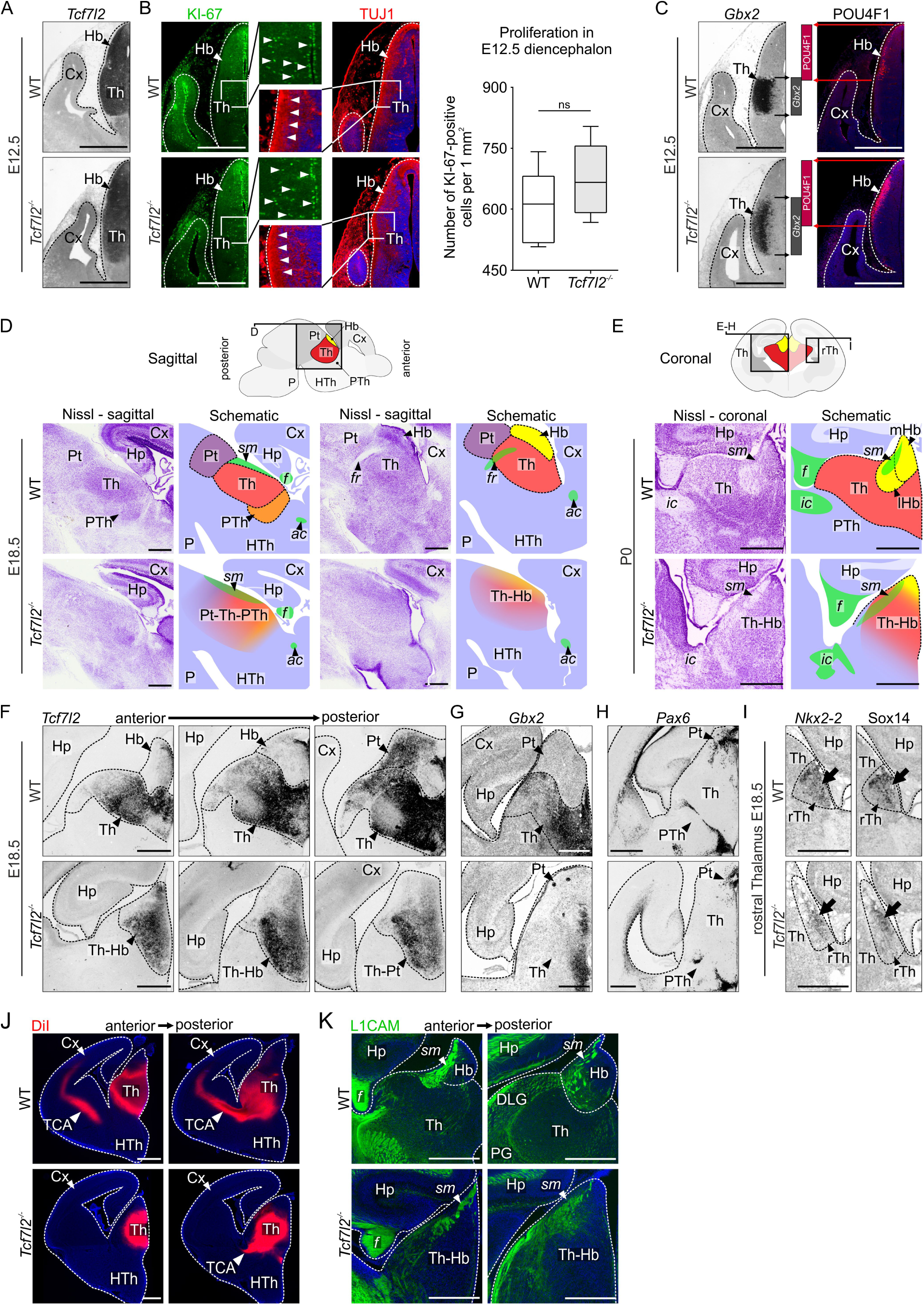
TCF7L2 controls the establishment of proper anatomy and axonal connections of the thalamo-habenular region in late gestation, but does not affect its early proliferation and neurogenesis. (A) *In situ* hybridisation staining with a *Tcf7l2* probe on E12.5 coronal brain sections. Expression from the *Tcf7l2* locus is maintained in the brain in *Tcf7l2^-/-^* embryos. **(B)** Immunofluorescent staining of the proliferation marker KI-67 antigen and neural marker TUJ1 (neuron-specific class III β-tubulin) on E12.5 coronal brain sections. Both in WT and *Tcf7l2*^-/-^ embryos, proliferating cells are visible along the ventricle, and young neurons are visible in the mantle zone (both marked with white arrowheads). Quantification of the number of proliferating cells in prosomere 2 revealed no significant changes between *Tcf7l2^-/-^* and WT embryos (WT n=3; *Tcf7l2^-/-^* n=3; *p*=0.0615). (C) *In situ* hybridisation staining with a *Gbx2* probe (early marker of the thalamus) and immunofluorescent staining of POU4F1 (early marker of the habenula) on E12.5 coronal brain sections. Both markers are expressed in *Tcf7l2^-/-^* embryos at this age, but the *Gbx2*-positive region and POU4F1-positive region spread into each other’s territory (emphasised by bars). Nissl staining in E18.5 sagittal **(D)** and coronal (**E**) brain sections. In *Tcf7l2^-/-^* embryos the shape of the thalamo-habenular region is changed and the anatomical boundaries are not visible. (F) *In situ* hybridisation staining with a *Tcf7l2* probe in consecutive E18.5 brain sections. Continued expression from the *Tcf7l2* locus indicates that neurons in the thalamo-habenular area were generated in prosomere 2. *In situ* hybridisation staining with *Gbx2* (partial marker of thalamus) **(G)** and *Pax6* (marker of pretectum and partial marker of prethalamus) **(H)** probes in E18.5 coronal brain sections. *Gbx2* expression in *Tcf7l2^-/-^* embryos becomes limited to the medial thalamus. (I) *In situ* hybridisation staining with *Nkx2-2* and *Sox14* probes (markers of rTh) on E18.5 brain sections. The expression of these markers is retained in *Tcf7l2^-/-^* embryos, although *Nkx2-2*- and *Sox14*-positive areas spread along the lateral wall of the thalamic region. **(J)** DiI tracing of thalamocortical tracts. There is little to no thalamocortical axon growth in *Tcf7l2^-/-^* embryos. **(K)** Immunofluorescent staining of the axonal marker L1CAM in the thalamo-habenular region. Topography of axonal fibers in thalamic and habenular areas is changed, *stria medullaris* is disorganised and less compact. ac, anterior commissure; Cx, cortex; DLG, dorsal lateral geniculate nucleus; f, fornix; Hb, habenula; Hp, hippocampus; HTh, hypothalamus; ic, internal capsule; lHb, lateral habenula; mHb, medial habenula; P, pons; PG, pregeniculate nucleus; Pt, pretectum; PTh, prethalamus; rTh, rostral thalamus; sm, stria medullaris; TCA, thalamocortical axons; Th, thalamus; Th-Hb, thalamo-habenular region. Scale bars represent 0.5 mm.

To investigate whether TCF7L2 is required for the initial acquisition of postmitotic molecular identities in the thalamus and habenula, we examined the expression of the *Gbx2* gene and POU4F1 protein (the earliest markers of postmitotic neurons in the thalamus and habenula, respectively) during neurogenesis. Both *Gbx2* mRNA and POU4F1 were highly expressed in the prosomere 2 area in WT and *Tcf7l2^-/-^* embryos on E12.5, indicating that their expression is not induced by TCF7L2. However, *Gbx2*- and POU4F1-positive areas expanded extensively into each other’s territory (Fig. 3C), suggesting a defect in thalamo-habenular boundary formation.

To further investigate if TCF7L2 regulates structural maturation of the prosomere 2 area, we focused on late gestation, when prosomere 2 neurons are already clustered into discrete nuclei and axonal connections are well-developed. The anatomy of the region was firstly analysed on E18.5 by Nissl staining. The boundaries between prosomere 2 and neighbouring structures, i.e., the prethalamus (rostral boundary) and pretectum (caudal boundary), were not morphologically detected in sagittal sections from *Tcf7l2^-/-^* embryos (Fig. 3D). On coronal sections, the whole region was reduced in the radial dimension, resulting in an oval-like shape, the habenula was fused with the thalamus, and nuclear groups within prosomere 2 were not clearly demarcated (Fig. 3E). To identify prosomere 2 and its main subdivisions, we analysed the expression pattern of several diencephalon markers by *in situ* hybridization. We used a *Tcf7l2* probe to stain prosomere 2 and pretectum (prosomere 1) (Fig. 3F), *Gbx2* probe to stain the thalamus (Fig. 3G), *Pax6* probe to stain the prethalamus (prosomere 3) and pretectum (Fig. 3H), and *Nkx2-2* and *Sox14* probes to stain the rTh (Fig. 3I). These stainings confirmed that although thalamic and habenular neurons were generated, the thalamo-habenular area was fused and malformed in *Tcf7l2^-/-^* embryos. Furthermore, the boundary between prosomere 2 and 3 that was demarcated by *Pax6* staining was disrupted. In addition, we noticed that *Gbx2* staining was still present in the periventricular area, where the youngest neurons are located, but absent in the intermediate and superficial portions of the thalamus (Fig. 3G), suggesting premature downregulation of *Gbx2* in the mutant embryos.

Then, we paid attention to the connectivity of the thalamo-habenular region. Consistent with a previous report (Lee et al., 2017), the major habenular efferent tract, i.e., the fasciculus retroflexus (habenulo-interpeduncular tract), was virtually absent in the Nissl-stained KO brains (Fig. 3D) and thalamocortical axons were not detected by DiI tracing (Fig. 3J). We also noticed that the bundles of stria medullaris, which include afferent fibres from the basal forebrain and lateral hypothalamus to the habenula, and were previously reported to be normal in *Tcf7l2^-/-^* embryos on E16.5 (Lee et al., 2017), were less compact on E18.5 (Fig. 3E). To observe the formation of these connections, we immunostained brain sections with an antibody specific for L1 cell adhesion molecule (L1CAM) which marks growing axons. This staining confirmed that stria medullaris were disorganised in E18.5 *Tcf7l2* KO embryos (Fig. 3K). It also showed major disruption in the general topography of axonal connections in the whole *Tcf7l2^-/-^* thalamic area. This demonstrated that TCF7L2 is critically involved in the development of habenular and thalamic connectivity.

### Impaired cell clustering in the diencephalon

Because anatomical boundaries were blurred and afferent connections were topographically disorganised in prosomere 2, we hypothesized that cells in this region do not cluster properly in *Tcf7l2^-/-^* embryos. To examine prosomere 2 internal and external boundaries at the cellular level, we stained brain sections with ani-PAX6, anti-NKX2-2 and anti-POU4F1 antibodies. Prosomere 2 cells were identified with anti-TCF7L2 (WT mice) or anti-β-galactosidase (KO mice) antibodies. In control embryos, the thalamic area was delineated rostro-ventrally by a narrow strip of PAX6-positive cells in a prethalamic subdomain (Fig. 4A), which separated the thalamus from SIX3-positive prethalamic area (Fig. 4B), and NKX2-2-positive area of the ventral lateral geniculate nucleus and intergeniculate leaflet, which derive from the rTh, did not overlap with TCF7L2-high area of the caudal thalamus (Fig. 4C). Habenular cells were easily identified by POU4F1 staining in WT embryos, and the differences in cell densities allowed discernment of its lateral and medial parts (Fig. 4D). In contrast, in *Tcf7l2* KO embryos, many PAX6-positive, NKX2-2-positive and POU4F1-positive cells were intermingled into the neighbouring thalamic territories (Fig. 4A-D). Consequently, the border between prosomere 2 and 3 was devoid of its sharpness (Fig. 4A-B), and thalamo-habenular border did not exist (Fig. 4D). This demonstrated a critical role of TCF7L2 in the segregation of cells in subregional clusters in prosomere 2.

**Figure 4.**
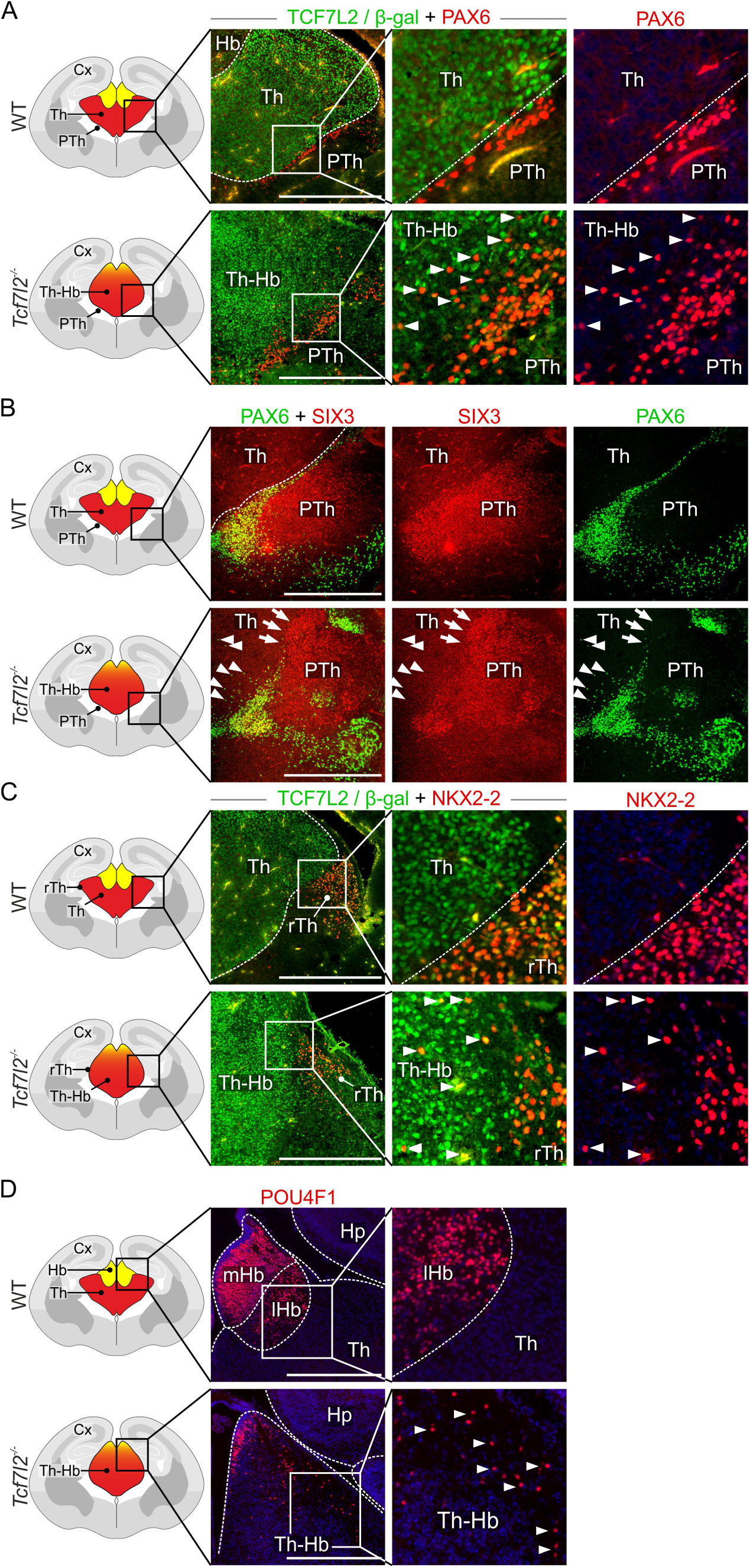
TCF7L2 controls the establishment of anatomical borders in the thalamus and habenula. (A) Immunofluorescent staining of TCF7L2 (in WT mice) or β-galactosidase (in *Tcf7l2^-/-^* mice) and co-staining of PAX6 in E18.5 coronal brain sections. PAX6-positive cells from prethalamus abnormally invade the thalamus in *Tcf7l2^-/-^* embryos (marked with white arrowheads). **(B)** Immunofluorescent staining of PAX6 and SIX3 (markers of prethalamic subregions) in E18.5 coronal brain sections. In *Tcfl2^-/-^* embryos, SIX3-positive cells (marked with white arrows) and PAX6-positive cells (marked with white arrowheads) intermingle into the thalamic region. **(C)** Immunofluorescent staining of TCF7L2 (in WT mice) or β-galactosidase (in *Tcf7l2^-/-^* mice) and co-staining of NKX2-2 in E18.5 coronal brain sections. NKX2-2-positive cells from rostral thalamus (marked with white arrowheads) abnormally invade the thalamus in *Tcf7l2^-/-^* embryos. **(D)** Immunofluorescent staining of the habenular marker POU4F1 in coronal brain sections. The number of POU4F1-positive cells is reduced in the habenular region in *Tcf7l2^-/-^* embryos. The remaining POU4F1-positive cells either accumulate in the subventricular region of the habenula or spread into the thalamic area (marked with white arrowheads). Cx, cortex; Hb, habenula; lHb, lateral habenula; mHb, medial habenula; Pt, pretectum; PTh, prethalamus; rTh, rostral thalamus; sm, stria medullaris; Th, thalamus; Th-Hb, thalamo-habenular region. Scale bars represent 0.5 mm.

### Disrupted prosomere 2-specific regulatory network and altered expression of morphogenesis effector genes

To investigate the possible role of TCF7L2 in regulating a genetic program of region-specific maturation program in prosomere 2, we analysed global gene expression by RNA-seq in prosomere 2 in WT and *Tcf7l2*^-/-^ embryos on E18.5. 210 genes were significantly downregulated and 113 were upregulated in KO embryos by ≥ 0.4 or ≤ −0.4 log_2_ FC (Fig. 5A, Table S1). Gene ontology (GO) term analysis of the differentially expressed genes (DEGs) revealed an overrepresentation of genes that are involved in transcription factor activity, anatomical structure development, neuron differentiation, axon guidance, cell adhesion, regulation of cell migration, regulation of transcription and synaptic signalling (Fig. 5A, Table S2). To determine whether the E18.5 DEGs from these groups are specific for prosomere 2, we inspected the corresponding *in situ* hybridisation images of brain sections in the Allen Brain Atlas (ABA). 100% of the downregulated genes in the selected groups were enriched in the thalamus, habenula, or both (Fig. 5A). Among them were prosomere 2-specific transcription factor genes, including known regulators of thalamic or habenular development - *Rora*, *Foxp2* and *Nr4a2* (Ebisu et al., 2016; Quina et al., 2009; Vitalis et al., 2017), cell adhesion molecule genes such as *Cdh6* (Yuge et al., 2011) and axon guidance genes such as *Epha4*, *Ntng1*, and *Robo3* that encode important regulators of the guidance of thalamic or habenular efferent connections and the segregation of neurons in this region (Belle et al., 2014; Braisted et al., 2000; Dufour et al., 2003; Lehigh et al., 2013). Also excitability genes, such as thalamus-enriched serotonin transporter gene *Slc6a4* that is expressed only during embryogenesis (Chen et al., 2015) were downregulated in *Tcf7l2* KO embryos. In contrast, the list of the upregulated genes was dominated by ones that were specifically expressed along thalamic borders, for example *Reln* that is involved in neuronal migration and positioning (Hirota and Nakajima, 2017).

**Figure 5.**
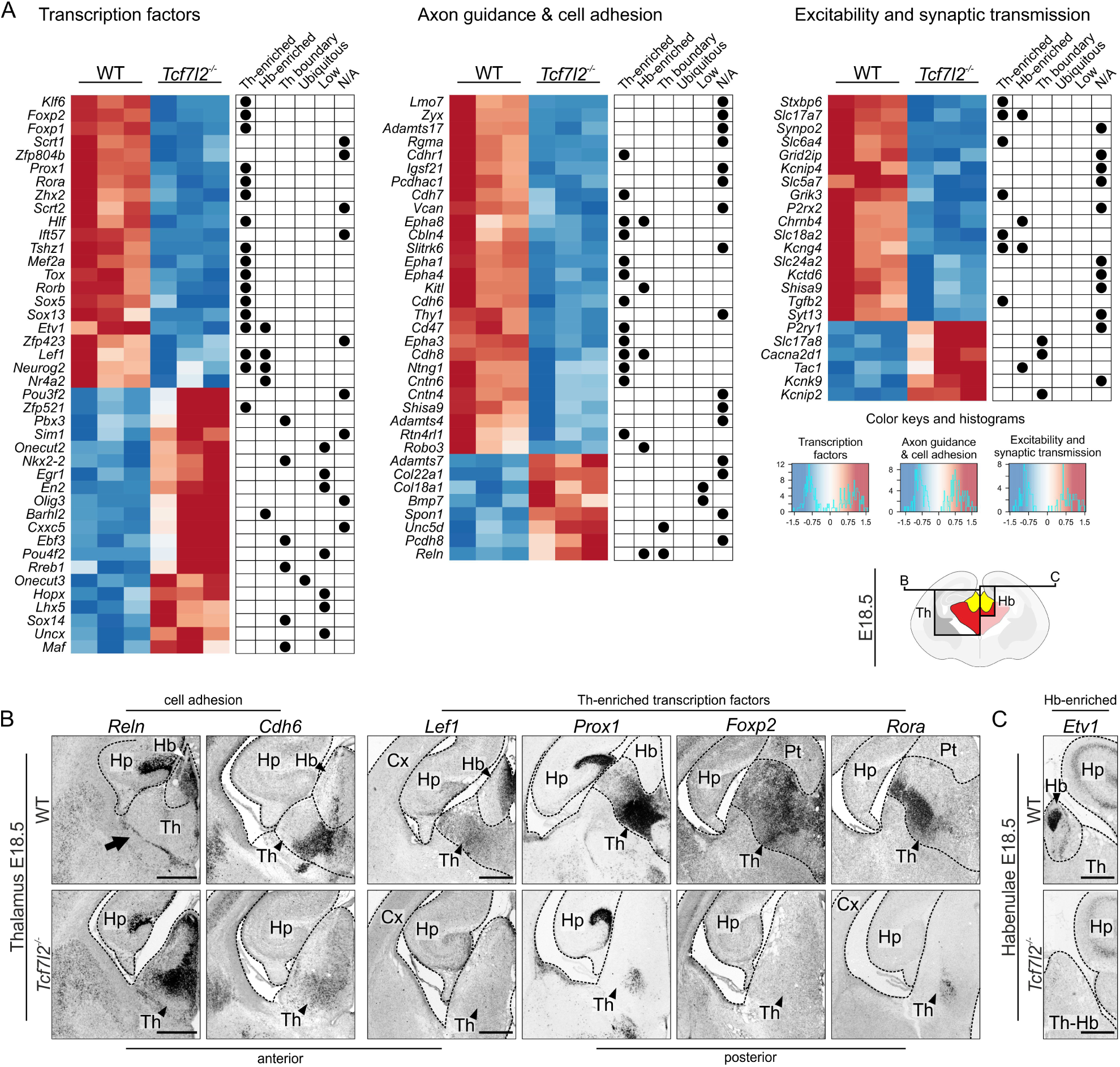
TCF7L2 orchestrates genetic program of morphological maturation in the thalamo-habenular region. (A) Hierarchical clustering of differential gene expression in prosomere 2 in *Tcf7l2*^-/-^ embryos on E18.5. Heatmaps of log2 expression fold changes in the DEGs from overrepresented groups of: (i) transcription factors; (ii) axon guidance and cell adhesion molecules; (iii) genes related with neuron excitability and synaptic transmission, and their localisation in the brain (assessed in the Allen Brain Atlas, marked with black dots to the right of the matrices). The knockout of *Tcf7l2* reduces the expression of most thalamus- and habenula-specific developmental genes. Th-enriched, genes with expression enriched in the thalamus; Hb-enriched, genes with expression enriched in the habenula; Th boundary, genes expressed at the boundary of prosomere 2; Ubiquitous, genes ubiquitously expressed throughout the brain; N/A, genes whose spatial expression is not available at Allen Brain Atlas **(B)** *In situ* hybridisation staining with *Reln* (marker of habenula and prethalamus) and *Cdh6*, *Lef1, Prox1, Foxp2* and *Rora* probes (markers of thalamic subregions) in E18.5 coronal brain sections. The expression of thalamic markers is either abolished, or drastically reduced in *Tcf7l2^-/-^* embryos at this age. Conversely, *Reln* expression becomes abundant in the mutant thalamus. **(C)** *In situ* hybridisation staining with *Etv1* probe (marker of the habenula) in E18.5 brain sections. The expression of this marker is abolished in *Tcf7l2^-/-^* embryos at this age. Cx, cortex; Hb, habenula; Hp, hippocampus; PTh, prethalamus; rTh, rostral Thalamus; Th, thalamus; Th-Hb – thalamo-habenular region. Scale bars represent 0.5 mm.

To confirm that the knockout of *Tcf7l2* caused a misexpression of prosomere 2-enriched or depleted genes, we validated several of the identified E18.5 DEGs by *in situ* hybridisation. We observed strong ectopic expression of *Reln* and decreased expression of *Cdh6* in the thalamus in the mutant embryos (Fig. 5B). The expression of transcription factor genes *Rora*, *Lef1, Foxp2*, and *Prox1*, which are subregional thalamic markers, was virtually absent in the thalamic area in *Tcf7l2^-/-^* embryos (Fig. 5B). Furthermore, *Tcf7l2* knockout abolished the expression of habenular markers *Lef1* (Fig. 5B) and *Etv1* (Fig. 5C) in different habenular subregions.

These results revealed that TCF7L2 controls a network of prosomere 2 subregional transcription factors and regulates region-specific axon guidance and cell migration related genes.

### Normal acquisition of glutamatergic identity but impaired expression of postnatally induced synaptic and excitability genes in the thalamus

We then asked if TCF7L2 acts also as a terminal selector of thalamic phenotype, which is underlined by the expression of region-specific genes that determine neurotransmitter identity and electrophysiological characteristics. Habenular and thalamic neurons in rodents (except for rTh-derived GABAergic interneurons (Evangelio et al., 2018)) are glutamatergic and express high levels of a vesicular glutamate transporter VGLUT2 (Fremeau et al., 2001; Herzog et al., 2001). To determine whether TCF7L2 is involved in the adoption of glutamatergic fate in the thalamus and habenula, we examined the expression patterns of *Vglut2/Slc17a6* and *Gad67/Gad1* (marker of GABAergic neurons) in the diencephalon in *Tcf7l2^-/-^* embryos and *Cck^Cre^:Tcf7l2^fl/fl^* P60 adult mice. Both knockout strains exhibited a pattern of GABAergic and glutamatergic cell distribution that was similar to the wild-type condition, with predominant *Vglut2/Slc17a6* expression in prosomere 2 (Fig. 6A). Thus, TCF7L2 is not involved in the specification and maintenance of VGLUT2-identity in prosomere 2.

**Figure 6.**
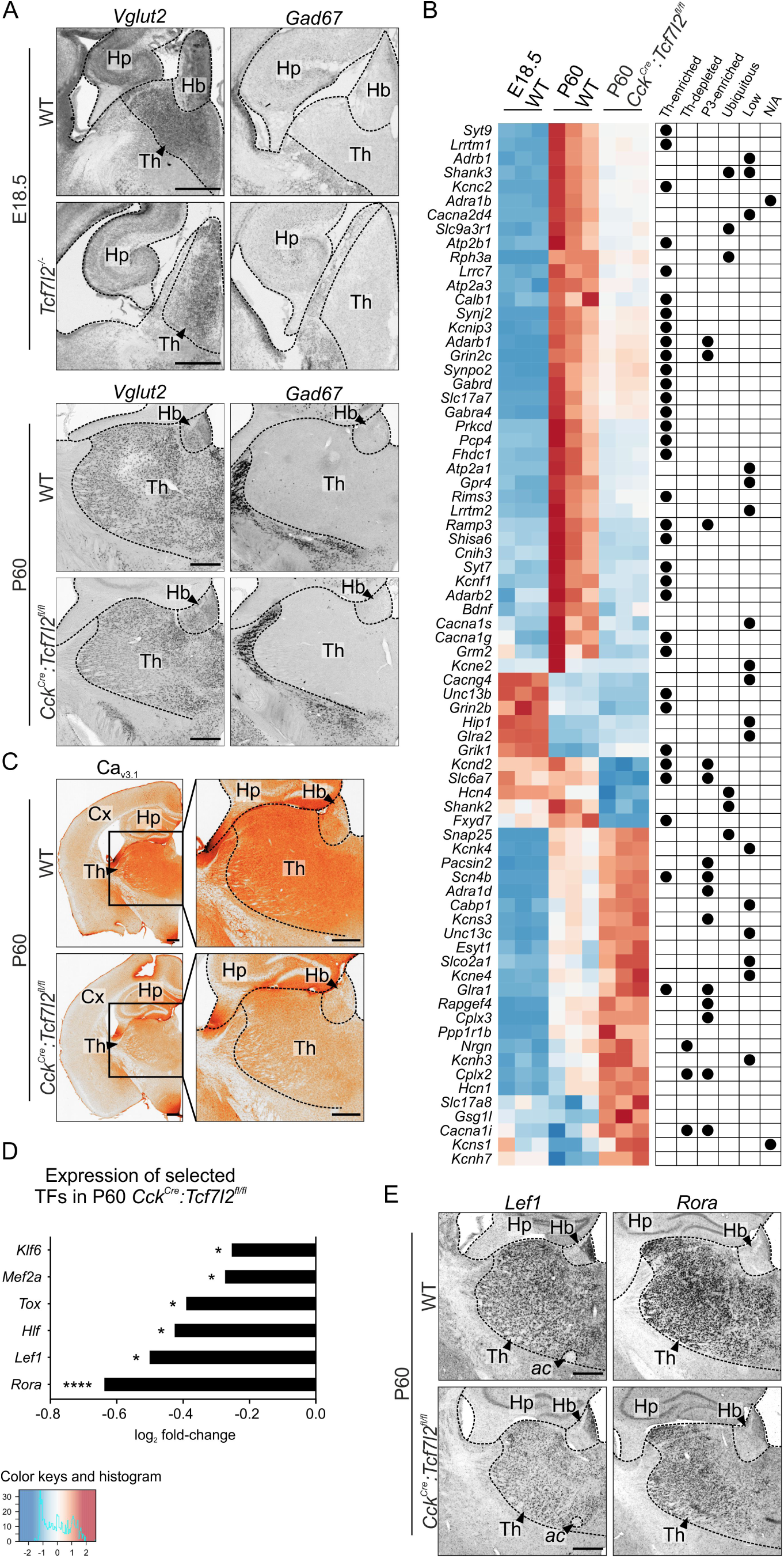
TCF7L2 controls the expression of terminal excitability genes, but not VGLUT2 identity in the thalamus. (A) *In situ* hybridisation staining with *Vglut2/Slc17a6* and *Gad1/Gad67* probes in E18.5 and P60 coronal brain sections. Both in E18.5 *Tcf7l2^-/-^* and P60 *Cck^Cre^:Tcf7l2^fl/fl^* mice the thalamic area retains predominant glutamatergic identity. **(B)** Hierarchical clustering of differential gene expression in prosomere 2 in *Cck^Cre^:Tcf7l2^fl/fl^* mice on P60 compared to control mice on E18.5 and P60. Heatmap of log2 expression fold changes in the DEGs from overrepresented groups of: (i) ion-channels; (ii) neurotransmitter receptors and transmitters; (iii) G-proteins and synaptic vesicle proteins, and their localisation in the brain (assessed in the Allen Brain Atlas, marked with black dots to the right of the matrices). Postnatal loss of TCF7L2 reduces the expression of thalamus-enriched excitability genes. Th-enriched, genes with expression enriched in the thalamus; Th-depleted, genes that are expressed ubiquitously in the brain except for the thalamus; P3-enriched genes with expression enriched in prosomere 3; Ubiquitous, genes ubiquitously expressed throughout the brain; Low, genes with overall low expression throughout the brain; N/A, genes whose spatial expression is not available at Allen Brain Atlas. **(C)** DAB immunohistochemical staining of Ca_v3.1_ Ca^2+^ ion channel (encoded by *Cacna1g* gene) in P60 coronal brain sections. The level of Cav_3.1_ protein is lower in *Cck^Cre^:Tcf7l2^fl/fl^* thalamus when compared to control. **(D)** Bar plot of log_2_ transformed transcript levels of differentially expressed transcription factor genes in P60 *Cck^Cre^:Tcf7l2^fl/fl^* compared to control (p≤0.05=*; p≤0.0001=****). **(E)** *In situ* hybridisation staining with *Rora* and *Lef1* probes (markers of thalamic subregions) in P60 coronal brain sections. In *Cck^Cre^:Tcf7l2^fl/fl^* mice the expression of *Rora* is significantly lower in the medial part of the thalamus when compared to control, while the expression of *Lef1* is lower throughout the whole thalamus. ac, anterior commissure; Hb, habenula; Hp, hippocampus; Th, thalamus. Scale bars represent 0.5 mm.

To investigate the hypothesis that TCF7L2 regulates terminal gene batteries, we used *Cck*^Cre^:*Tcf7l2*^fl/fl^ mice with the postnatal knockout of *Tcf7l2* and compared global gene expression profiles in the thalamus between these mutant mice and WT mice on P60 by RNA-seq. 310 genes were significantly downregulated and 229 were upregulated in KO mice by ≥ 0.4 or ≤ −0.4 log_2_ FC (Fig. 6B, Table S3). GO term enrichment analysis of the P60 DEGs revealed significant enrichment with terms that clustered into groups of synaptic proteins and regulators of membrane conductance: regulation of ion transport, voltage-gated channel activity, regulation of membrane potential, G-protein coupled receptor signalling pathway, regulation of trans-synaptic signalling and regulation of synapse organisation (Fig. 6B, Table S4). To investigate if TCF7L2 regulates thalamus-specific or generic neuronal genes, we examined spatial expression profiles of the identified genes in this cluster in the ABA. Vast majority of the downregulated genes are expressed specifically thalamic (Fig. 6B), such as *Kcnc2* and *Cacna1g*, which encode subunits of K_v3.2_ voltage-gated potassium channels and Ca_v3.1_ voltage-gated calcium channels, respectively (Kasten et al., 2007; Kim et al., 2001). The downregulation of Ca_v3.1_ was further confirmed by immunohistochemistry (Fig. 6C). Also the habenular and thalamic glutamate transporter gene *Vglut1*/*Slc17a7* was downregulated, but not *Vglut2/Slc17a6* that encodes the main thalamic glutamate transporter, consistently with the *in situ* hybridisation results (Fig. 6A). Conversely, only a few of the upregulated excitability genes were thalamus-enriched.

To investigate the hypothesis that TCF7L2 regulates a genetic program of terminal selection that is activated postnatally, we crossed the selected group of genes with the list of genes that were differentially expressed between E18.5 and P60 in WT mice. Almost 90% of the synaptic/excitability P60 DEGs that were thalamus-enriched were induced after embryogenesis, confirming that TCF7L2 functions as a terminal selector during postnatal development.

Transcription factor genes were not overrepresented in the P60 DEGs. However, 6 out of 15 thalamus-enriched subregional transcription factor genes, which were downregulated in *Tcf7l2*^-/-^ embryos, were also significantly downregulated in *Cck*^Cre^:*Tcf7l2*^fl/fl^ mice on P60 by ≥ 0.4 or ≤ −0.4 log_2_ FC (Fig. 6D), including *Rora* and *Lef1*, confirmed by *in situ* hybridization (Fig. 6E).

Thus, postnatally, TCF7L2 maintains thalamic regulatory network and regulates a battery of genes that shape terminal electrophysiological identities of thalamic neurons.

### Direct regulation of thalamic terminal effector genes by TCF7L2

To understand how TCF7L2 regulates terminal effector genes in the thalamus, we performed ChIP-Seq analysis on the thalami isolated from adult WT mice. Analysis resulted in 4625 peaks in anti-TCF7L2 precipitated samples with fold enrichment ≥10 over input sample, which annotated to 3791 unique genes (Table S5). We used MEME-ChIP suite to discover motifs enriched in TCF7L2 bound peaks. The most significantly enriched de novo motif was identical to TCF7L2/TCF7L1 consensus binding site (Fig. 7A), proving that most of TCF7L2 binding in the adult thalamus occurred through a classical TCF7L2 binding site.

**Figure 7.**
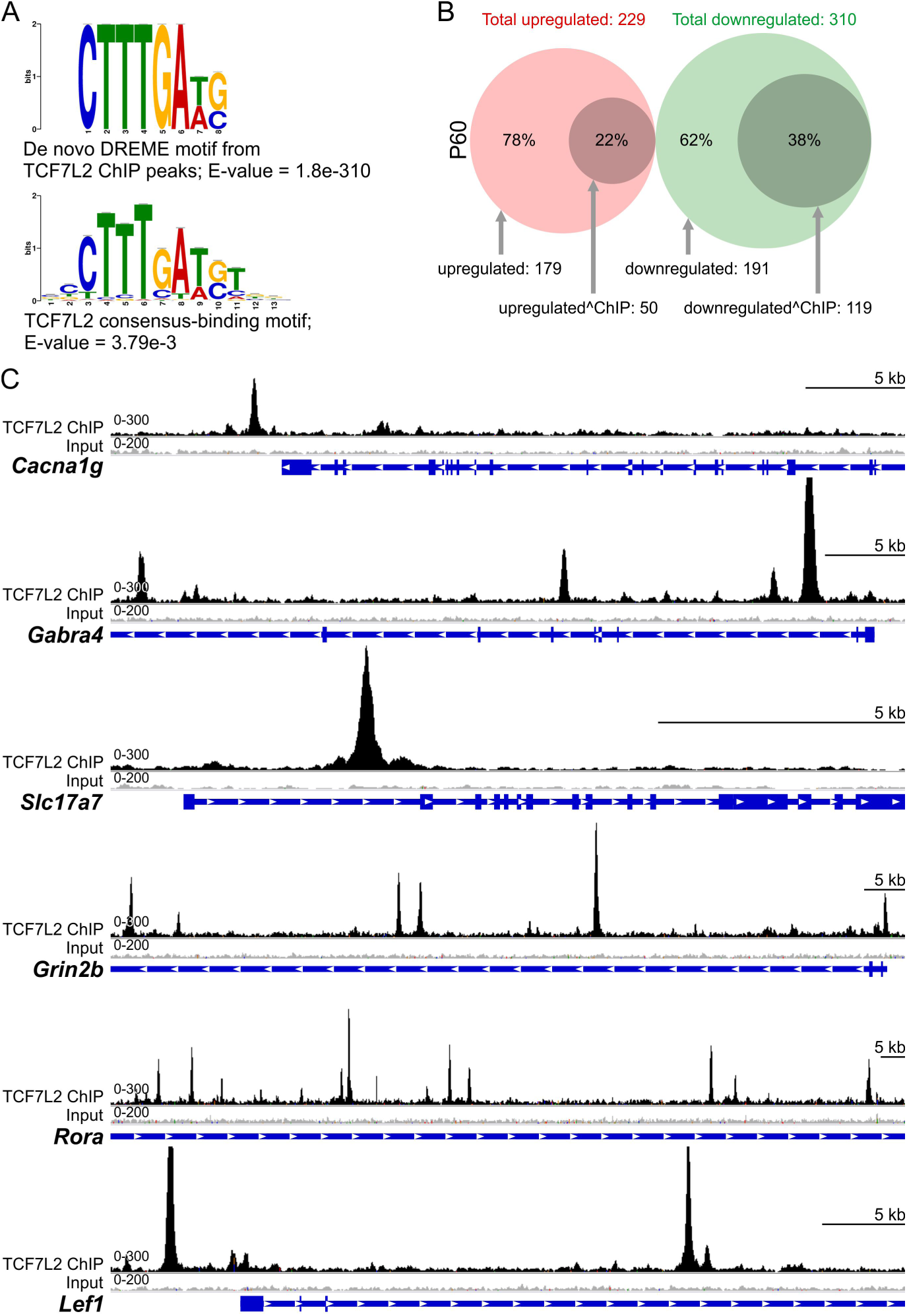
TCF7L2 directly regulates terminal effector genes in the thalamus. (A) De novo motif discovery analysis with MEME-ChIP: top: de novo DREME motif identified from TCF7L2 ChIP peaks (e-value = 1.8e-310); bottom: match of de novo motif to known TCF7L2 consensus binding site (e-value: = 3.79e-3). **(B)** Venn diagrams showing overlap between differentially expressed genes (DEGs) in P60 *Cck^Cre^:Tcf7l2^fl/fl^* thalami (RNA-Seq n=3) and genes bound by TCF7L2 in a P60 WT ChIP-Seq (n=2). Upregulated genes are marked in red, downregulated genes are marked in green, the DEGs that are positive for TCF7L2 peaks are marked with dark circles within both clusters. **(C)** TCF7L2 binding profiles (in black) in adult mouse thalami for *Cacna1g, Gabra4, Slc17a7, Grin2b, Rora, Lef1.* Signal from input is in grey.

GO term enrichment analysis of the genes significantly bound by TCF7L2 revealed an overrepresentation of genes related to membrane potential regulation and synaptic signalling, potassium ion transmembrane transport, and also regulation of neuron projection development and cell adhesion (Table S6). This showed that TCF7L2 regulates the expression of neural terminal effector genes. For example, TCF7L2 localised on genes that encode subunits of GABA receptors (e.g., thalamus-enriched*, Gabra4, Gabbr2, Gabrd*) and glutamate receptors (e.g., thalamus-enriched*, Grin2b, Grm1* and *Grm7*).

TCF7L2 binding was detected in 38% genes that were downregulated and 22% genes that were upregulated in *Cck^Cre^:Tcf7l2^fl/fl^* thalami on P60 (Fig. 7B, Table S5), suggesting that TCF7L2 acts directly as an activator rather than repressor. TCF7L2 binding sites were more abundant in the genes that were downregulated in the mutant mice (199 peaks on 119 genes in total), compared to the upregulated ones (64 peaks on 50 genes in total) (Fig. 7B-C), corroborating this conjecture. Among the downregulated loci with TCF7L2 binding peaks were many thalamus-enriched genes that encode proteins involved in synaptic signalling or adhesion, such as *Synpo2, Prkcd* and *Ramp3*, *Cacna1g*, *Kcnc2*, *Lrrtm1*, *Lingo1*, *Ntng1* and aforementioned *Gabra4*, *Gabrd* and *Grin2b*. This confirmed that TCF7L2 is directly involved in the regulation of genes that define thalamic neuron terminal identity. Furthermore, TCF7L2 binding sites in the *Rora* and *Lef1* loci revealed that TCF7L2 directly controls thalamic regulatory network.

### Severe impairments in excitability of thalamic neurons in adult mice

To functionally test the role of TCF7l2 in the regulation of thalamic neurons electrophysiological properties, we used whole-cell patch-clamp recordings in *Cck^Cre^:Tcf7l2^fl/fl^* mice from targeted thalamocortical neurons in the ventrobasal complex (VB). We first tested basic properties of VB neurons. Resting potential and capacitance were similar in WT and KO mice, but input resistance significantly decreased in KO mice (Fig. 8A). We then examined whether the targeted neurons were able to evoke action potentials (APs; e.g., tonic, burst, and rebound burst modes) that are typical for thalamocortical cells. We used depolarizing current steps to evoke both sustained trains of APs (tonic) and burst firing at the beginning of a train (Fig. 8B-C). Thalamic cells produced fewer APs in *Cck^Cre^:Tcf7l2^fl/fl^* mice in both tonic and burst firing modes (Fig. 8D). Rebound bursts, which are crucial for the response of thalamocortical neurons to inhibitory input, were evoked by steps of hyperpolarizing current (Fig. 8E). Most neurons from *Tcf7l2* KO mice did not show any rebound bursts at the hyperpolarizing membrane potential (−65 mV; Fig. 8F). Hyperpolarizing steps that were applied at the resting membrane potential (∼ −57 mV) evoked rebound burst spiking in VB neurons in *Cck^Cre^:Tcf7l2^fl/fl^* mice, but the number of spikes was approximately two-times lower compared to WT mice (Fig. 8G). These dramatic impairments in electrophysiological responses demonstrated that TCF7L2 is essential for the establishment of unique excitability and firing patterns in thalamocortical neurons.

**Figure 8.**
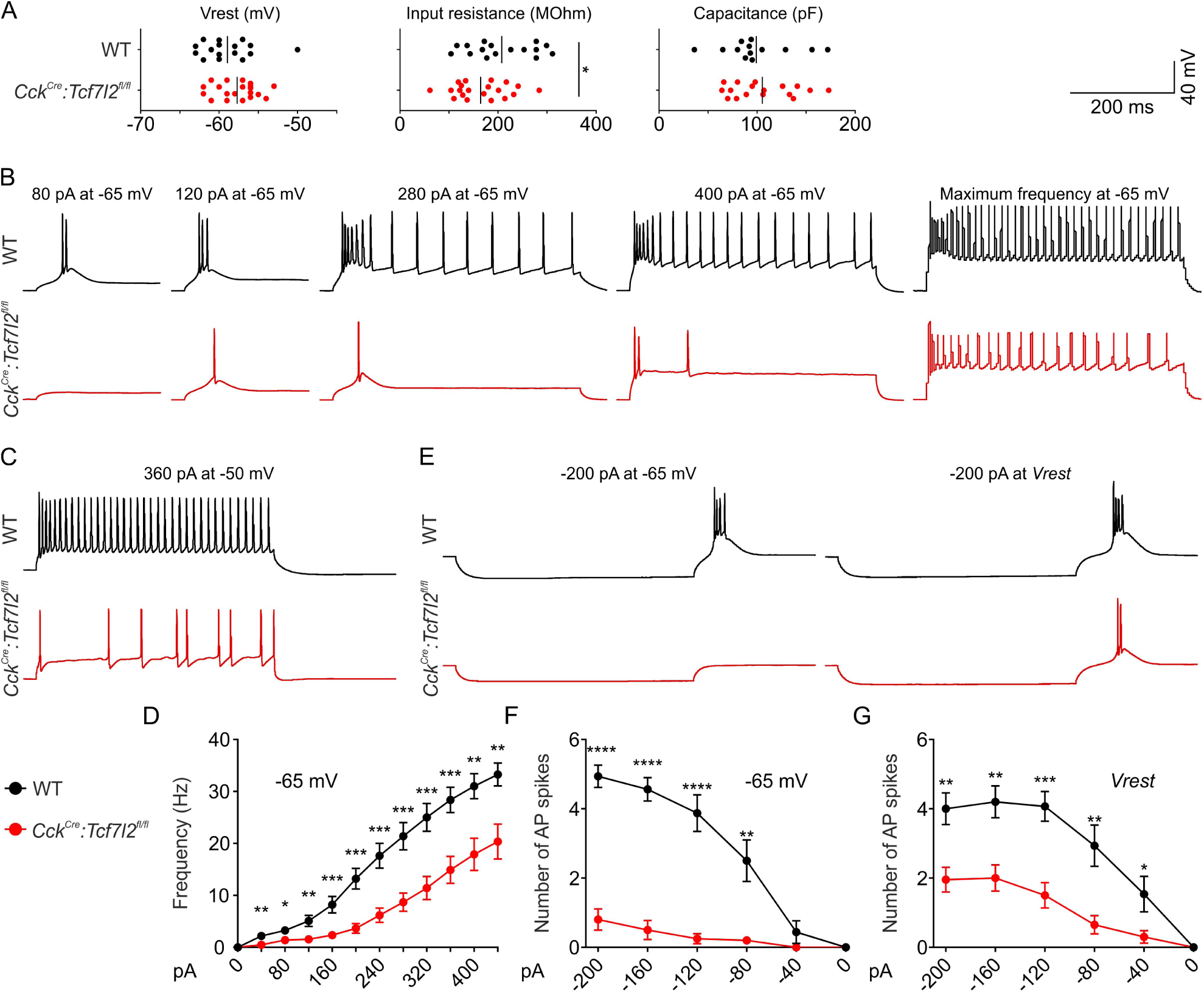
TCF7L2 is essential postnatally for the development of burst and tonic firing patterns in thalamocortical relay neurons. (A) Basic electrical properties of cell membrane measured in VB neurons in brain slices. Thalamocortical neurons in *Cck^Cre^:Tcf7l2^fl/fl^* mice have lower input resistance (WT n=16, *Cck^Cre^:Tcf7l2^fl/fl^* n=20, *p* value=0.0443), but their resting potential (WT n=16, *Cck^Cre^:Tcf7l2^fl/fl^* n=20, *p* value= 0.1112) and membrane capacitance (WT n=13, *Cck^Cre^:Tcf7l2^fl/fl^* n=17, *p* value=0.6725) are not significantly changed. Representative traces from whole-cell patch-clamp recordings from VB neurons in brain slices at −65 mV membrane potential and increasing depolarizing current inputs (burst and tonic spikes) **(B)**, and at −50 mV membrane potential and depolarizing current inputs (tonic spikes) **(C)**. **(D)** Frequency of spikes evoked by increasing depolarizing currents at −65 mV, as represented in (B). **(E)** Representative traces from whole-cell patch-clamp recordings from VB neurons in brain slices at −65 mV or the resting membrane potential (*Vrest*, ∼ −57 mV) and hyperpolarizing current inputs (rebound bursts of spikes). Number of spikes per a rebound burst at −65 mV **(F)** or at the resting membrane potential (*Vrest*) **(G)**, as represented in (E). In *Tcf7l2* mutant neurons, the frequency of spikes evoked at −65 mV membrane potential is much lower, and rebound bursts of spikes are virtually absent at this condition or are lower in number at the resting membrane potential. Tonic spikes are evoked at −50 mV membrane potential, but their frequency is significantly lower in *Tcf7l2* mutant neurons (WT n=15-16; *Cck^Cre^:Tcf7l2^fl/fl^*, n=18-20; p≤0.05=*; p≤0.01=**; p≤0.001=***; p≤0.0001=****; error bars indicate SEM).

## Discussion

Little is understood about the way in which the lengthy process of postmitotic differentiation is regulated in the vertebrate brain. The present study identifies TCF7L2 as a master regulator of regional transcription factors and a selector of stage-specific developmental programs that switch postnatally from morphological to electrophysiological maturation.

### TCF7L2 and a network of transcription factors regulate morphological maturation of the thalamus and habenula

TCF7L2 is the only developmentally regulated transcription factor that is expressed throughout prosomere 2 postmitotically (Nagalski et al., 2016). TCF7L2 is not critically involved in the induction of pan-thalamic and pan-habenular identities that are defined by *Gbx2* and *Pou4f1* expression, respectively, because the expression of these markers was not abolished in *Tcf7l2* KO embryos during neurogenesis (E12.5). Conversely, GBX2 does not activate *Tcf7l2* expression (Lee et al., 2017). However, the maintenance of *Gbx2* and *Pou4f1* expression may depend on TCF7L2 in maturating cells. *Gbx2* staining was absent in the intermediate and superficial portions of the mutant thalamus on E18.5 in *Tcf7l2* KO mice, and was present only in the periventricular area on E18.5, where the youngest neurons are located, suggesting premature downregulation of *Gbx2*. Similarly, POU4F1-positive cells accumulated in periventricular area, but were scarce outside of this zone in the mutant mice. A previous research suggested that TCF7L2 inhibits habenular identity (Lee et al., 2017). Our results do not support this hypothesis, because, in addition to *Gbx2* and *Pou4f2*, sub-habenular markers (e.g., *Etv1*) as well as sub-thalamic markers (e.g., *Foxp2, Lef1*, *Prox1* and *Rora*) were downregulated in E18.5 *Tcf7l2*^-/-^ embryos. *Rora* and *Lef1* proved to be direct targets of TCF7L2 in ChIP-seq assay, and our previous *in vitro* results showed that TCF7L2 can regulate promoters of thalamic factors *Gbx2*, *Pou4f1*, *Rora* and *Foxp2*, and habenular *Nr4a2* and *Etv1* (Nagalski et al., 2016).

Striking similarities between phenotypes in *Tcf7l2*^-/-^ and *Gbx2*^-/-^ embryos, such as impaired thalamic nucleogenesis and the absence of thalamocortical projections (Chatterjee et al., 2012; Chen et al., 2009), suggests that at least some TCF7L2 functions are mediated by GBX2. The expression of *Cdh6* (Cadherin 6) and *Ntng1* (Netrin G1) was disrupted in *Gbx2*^-/-^ as well as *Tcf7l2*^-/-^ embryos (Chen et al., 2009; Mallika et al., 2015), indicating that these genes might be regulated by TCF7L2 directly or in cooperation with GBX2. Less severe impairments in the establishment of thalamocortical connections were reported in *Foxp2* (Ebisu et al., 2016) and *Rora* (Vitalis et al., 2017) KO mice, and mice with either knockout exhibited molecular identity impairments that were restricted to subregions of the thalamus. These two genes were also downregulated in both *Tcf7l2*^-/-^ and *Gbx2*^-/-^ embryos. Similarities also exist in the habenula between *Tcf7l2^-/-^* and *Pou4f1*^-/-^ embryos. For example, in both cases, the expression of habenular *Rgma* (Repulsive guidance molecule A) and *Epha8* (Ephrin type-A receptor 8) decreased (Quina et al., 2009). However, anatomical impairments in the habenula were much more severe in *Tcf7l2^-/-^* embryos. Habenulo-interpeduncular tracts, which were affected but still present in *Pou4f1^-/-^* embryos (Quina et al., 2009; Serrano-Saiz et al., 2018), were missing in *Tcf7l2^-/-^* embryos. This indicates that TCF7L2 plays a critical role in the establishment of habenular connections to a large extend independently of POU4F1. These data point to the role of TCF7L2 as a master regulator of subregional regulatory networks during axon guidance and cell clustering in prosomere 2.

### TCF7L2 controls the acquisition of characteristic excitability patterns in the thalamus

Many genes that were downregulated in mice with the complete or postnatal *Tcf7l2* knockout and/or genes which were occupied by TCF7L2 in ChIP-seq assay encode thalamus-enriched neurotransmitter receptors, voltage-gated ion channels, and other proteins that are involved in neural signal transmission, consistently with our previous *in silico* predictions (Wisniewska et al., 2012).

Excitability and synaptic transmission parameters are specific to different classes of neurons and together ensure the proper functioning of neural circuits. Several thalamus-enriched excitability/synaptic genes whose expression increases during development were downregulated in adult *Tcf7l2* KO mice. Among these were *Cacna1g* and *Kcnc2*, which encode thalamus-enriched voltage-gated T-type Ca_v3.1_ calcium channels and K_v3.2_ potassium channels, respectively. We demonstrated that these genes are direct targets of TCF7L2 in the thalamus, consistent with previous *in vitro* results (Wisniewska et al., 2010). Functional experiments in the present study demonstrated that thalamocortical relay neurons lose their proper tonic and burst firing patterns in the absence of TCF7L2. The downregulation of *Cacna1g* and *Kcnc2* most likely contributed to this phenotype, given that *Cacna1g* knockout resulted in the absence of burst firing (Kim et al., 2001), whereas K_v3.2_ channel inhibition suppressed the firing rate in tonic mode (Kasten et al., 2007) in thalamocortical neurons, similar to *Tcf7l2* knockout. These results implicate TCF7L2 in the regulation of the expression of postnatal genes that control thalamic terminal excitability patterns.

Although majority of neurotransmission-related genes are active in mature neurons, early release of neurotransmitters and transiently expressed receptors during embryogenesis regulates the process of brain wiring (Andreae and Burrone, 2018). Several thalamus-enriched neurotransmission genes that are switched off postnatally were downregulated in *Tcf7l2^-/-^* embryos, e.g., *Slc6a4*, which encodes serotonin transporter 1 that regulates arborisation of thalamocortical axons (Chen et al., 2015). Notably, variants of *SLC6A4* are associated with autism spectrum disorder (Coutinho et al., 2004), and the transient inhibition of *Slc6a4* during development produced abnormal emotional behaviours in adult mice (Ansorge et al., 2004). Further research should investigate the possible contribution of TCF7L2-related impairments during this developmentally sensitive time window to aetiology of social and affective deficits.

### TCF7L2 is a terminal selector in the thalamus

TCF7L2 meets the criteria of a thalamic terminal selector: *(i)* its expression is induced during neurogenesis and maintained in thalamic neurons throughout life (Nagalski et al., 2013), *(ii)* it directly binds to thalamic terminal differentiation genes, *(iii)* and its activity is required for electrophysiological maturation of the thalamus. In addition, TCF7L2 orchestrates morphological maturation of the thalamo-habenular region during embryogenesis. This double function differentiate TCF7L2 from majority of other known terminal selectors, which play minor roles in cell migration or axon guidance (Hobert, 2016), for example habenular selector POU4F2 (Serrano-Saiz et al., 2018). Other examples of terminal selectors that regulate axon guidance are the glutamatergic selector of corticospinal neurons FEZF2 and serotoninergic selector PET1 (Donovan et al., 2019; Lodato et al., 2014).

TCF7L2 is essential for the postnatal induction of terminal functional properties of thalamic neurons. However, unlike classic terminal selectors, TCF7L2 does not control neurotransmitter identity, which in the thalamus depends on VGLUT2. This implies that in the case of glutamatergic neurons in the thalamus, the selection of different terminal features is uncoupled. It is known form the studies of invertebrate and vertebrate neurons that the adoption of neurotransmitter identity and other terminal features are often linked (Serrano-Saiz et al., 2013). For example, in contrast to thalamic cells and TCF7L2, PET1 controls the early postmitotic adoption of neurotransmitter phenotype and the postnatal acquisition of excitability features by serotonergic cells in the raphe nucleus (Hendricks et al., 2003; Liu et al., 2010; Wyler et al., 2016), and neurotransmitter metabolism genes together with ion channel genes constitute a co-varying module in midbrain dopaminergic neurons (Tapia et al., 2018). Furthermore, studies on glutamatergic neurons in mice concluded that POU4F1 and FEZF2 are selectors of VGLUT1 identity in the medial habenula and corticospinal neurons, respectively, as well as regulators of many other neuron subtype–specific genes (Chen et al., 2008; Lodato et al., 2014; Serrano-Saiz et al., 2018). There are only few examples of separate regulation of neurotransmitter identity and other terminal features. In *C. elegans*, *unc-3* regulates cholinergic identity in command interneurons, but not other terminal features (Pereira et al., 2015), and *ttx-3* controls the terminal differentiation of neurosecretory-motor neurons, but not their serotoninergic identity (Zhang et al., 2014). More research is needed to build models of the postmitotic regulation of neurons in vertebrates and compare regulatory strategies between vertebrates and invertebrates.

## Conclusion

The present study sheds new light on vertebrate regulatory strategies in the postmitotic differentiation of molecularly diverse neurons that share glutamatergic identity. We found that temporarily separated developmental stages can be controlled by the same regional transcription factors. We provided evidence that a master regional transcription factor can regulate the molecular diversification of neurons within the region. Finally, we showed that electrophysiological maturation can be uncoupled from the selection of neurotransmitter identity. Considering that *Tcf7l2* is associated with mental disorders, our findings provided new insight into the aetiology of thalamic and habenular dysfunction that is observed in these disorders.

## Materials and Methods

### Animals

We used C57BL/6NTac-*Tcf7l2*^tm1a^(EUCOMM)Wtsi/WtsiIeg (*Tcf7l2*^tm1a^) mouse strain (Skarnes et al., 2011), with a trap cassette upstream of the critical exon 6 of the *Tcf7l2* gene. To generate *Cck*^Cre^:*Tcf7l2*^fl/fl^ strain, in which the knockout of *Tcf7l2* is induced in the thalamic area perinatally (Allen Brain Atlas, 2011), *Tcf7l2*^tm1a/+^ animals were first crossed with flippase-expressing mice (JAX stock #009086; (Farley et al., 2000)), and then with *Cck*^tm1.1(cre)Zjh^/J mice (*Cck*^Cre^:*Tcf7l2*^+/+^, JAX stock #012706; (Taniguchi et al., 2011)), which express Cre recombinase from the *Cck* promoter. *Cck^Cre^:Tcf7l2^+/+^* animals were used as wild-types. Mice were maintained on a 12 h/12 h light/dark cycle with *ad libitum* access to food and water. For the experimental procedures all mice were selected by PCR-based genotyping: Tcf7l2^tm1a^ & Tcf7l2^fl^ alleles: tcf_F – GGAGAGAGACGGGGTTTGTG; tcf_R – CCCACCTTTGAATGGGAGAC; floxed_PNF – ATCCGGGGGTACCGCGTCGAG; Tm1c_R – CCGCCTACTGCGACTATAGAGA; Cck^Cre^ allele: 11214 – GAGGGGTCGTATGTGTGGTT; 11215 – GGGAGGCAGATAGGATCACA; 9989 – TGGTTTGTCCAAACTCATCAA. All of the experimental procedures were conducted in compliance with the current normative standards of the European Community (86/609/EEC) and Polish Government (Dz.U. 2015 poz. 266).

### Brain fixation

*Tcf7l2^-/-^* and control mice were collected on embryonic day 12.5 (E12.5) or E18.5. Noon on the day of appearance of the vaginal plug was considered E0.5. Timed-pregnant dams were sacrificed by cervical dislocation, the embryos were removed and decapitated. E18.5 brains were dissected out and fixed overnight in 4% paraformaldehyde (PFA; Cat. no. P6148, Sigma-Aldrich) in 0.1 M phosphate-buffered saline (PBS; pH 7.4; Cat. no. PBS404, BioShop) at 4°C. E12.5 heads were fixed whole. *Cck*^Cre^:*Tcf7l2*^fl/fl^ and *Cck*^Cre^:*Tcf7l2*^+/+^ mice were sacrificed on P60-P75 (further referred to as P60) by pentobarbital sedation and perfusion with PBS and 4% PFA. Their brains were dissected out and fixed overnight in 4% PFA. For cryostat sections, the brains were sequentially transferred into 15% and 30% sucrose in PBS at 4°C until they sank. Next, E12.5 and E18.5 tissues were embedded in 10% gelatine/10% sucrose solution in PBS (Ferran et al., 2015a; Ferran et al., 2015b). Postnatal brains were transferred to O.C.T (Cat. no. 4583, Sakura Tissue-Tek). Tissues were frozen in −60°C isopentane. Sections (20 μm – embryos, 40 μm - adult) were obtained using a Leica CM1860 cryostat. Embryonic sections were mounted directly on Superfrost-plus slides (Cat. no. J1800AMNZ, Menzel-Gläser). Adult tissue was collected as free-floating sections into an anti-freeze solution (30% sucrose/30% glycerol in PBS). For DiI axon tracing, E18.5 immersion-fixed brains were kept in 4% PFA at 4°C. Stained sections were visualized under a Nikon Eclipse Ni-U microscope.

### Nissl staining

Brain sections were dehydrated in a series of ethanol solutions (50%, 70%, 95%, and 99.8%), cleared in xylene and rehydrated. The sections were rinsed in tap water and stained with 0.13% (w/v) Cresyl violet solution (Cat. no. CS202101, Millipore) for 4 min, rinsed and dehydrated again as described above. The slices were washed in xylene and mounted using EuKitt.

### In situ hybridisation

The hybridisation was performed using digoxigenin-UTP-labelled antisense riboprobes (sense & antisense), synthesized with the DIG RNA labelling Kit (Cat. no. 11175025910, Roche). Plasmids for synthesis of *Cdh6*, *Foxp2*, *Gad1/Gad67*, *Gbx2*, *Lef1*, *Prox1, Rora,, Tcf7l2* and *Vglut2/Slc17a6* probes come from the collection of José Luis Ferran; the other plasmids were kind gifts from: James Li from the Department of Genetics and Developmental Biology, University of Connecticut Health Center, Farmington, CT, USA (*Etv1*) (Chatterjee et al., 2014); Seth Blackshaw from the Johns Hopkins University School of Medicine, Baltimore, MD, USA (*Nkx2-2* and *Sox14*) (Shimogori et al., 2010); Tomomi Shimogori from the RIKEN Center for Brain Science, RIKEN, Saitama, Japan (*Reln*) (Chiara et al., 2012); David Price from the Centre for Integrative Physiology, University of Edinburgh, UK (*Pax6*) (Walther and Gruss, 1991). Chromogenic *in situ* hybridisation of cryosections was performed as described elsewhere (Ferran et al., 2015b; Puelles et al., 2016). The stained sections were washed in xylene and mounted using EuKitt (Cat. no. 03989, Sigma-Aldrich). No specific signal was obtained with sense probes (data not shown).

### Fluorescent immunohistochemistry

Frozen sections were washed in PBS with 0.2% Triton X-100 (PBST) and blocked with 5% normal donkey serum (NDS) for 1 h. The slides were then incubated with primary antibodies mixed in 1% NDS overnight at 4°C. Antibodies against TCF7L2 (1:500; Cat. no. 2569, Cell Signaling), Ca_v3.1_ (1:500; Cat. no. MABN464, Sigma-Aldrich, NeuroMab clone N178A/9), β-galactosidase (1:100; Cat. no. AB986, Merck Millipore), L1CAM (1:500; Cat. no. MAB5272, Merck Millipore), PAX6 (1:100; Cat. no. PRB-278P, Biolegend), KI-67 (1:100; Cat. no. AB9260, Merck Millipore), TUJ1 (1:65; Cat. no. MAB1637, Merck Millipore), NKX2-2 (1:50; Cat. no. 74.5A5, DSHB), SIX3 (1:100; Cat. no. 200-201-A26S, Rockland), POU4F1 (1:300; (Fedtsova and Turner, 1995)) were used. Sections were then incubated for 1 h with appropriate secondary antibody conjugated with Alexa Fluor 488 or 594 (1:500; Cat. no. A-21202, A-21207, and A-11076, ThermoFisher Scientific). The slides were additionally stained with Hoechst 33342 (1:10000; Cat. no. 62249, ThermoFisher Scientific), washed and mounted with Vectashield Antifade Mounting Medium (Cat. no. H1000, Vector Laboratories).

### DAB immunohistochemistry

Free-floating sections were washed in PBST, incubated in 0.3% H_2_O_2_ for 10 min, blocked with 3% normal goat serum (NGS) for 1 h and incubated with primary antibodies against TCF7L2 (1:1000) or POU4F1 (1:600) in 1% NGS overnight at 4°C. Next, sections were incubated for 1 h with biotinylated goat anti-rabbit antibody (Cat. no. BA-1000, Vector Laboratories) in 1% NGS, and then for 1 h in Vectastain ABC reagent (Cat. no. PK-6100, Vector Laboratories). Staining was developed using 0.05% DAB (Cat. no. D12384, Sigma-Aldrich) and 0.01% H_2_O_2_. Next, sections were mounted onto Superfrost plus slides, washed in xylene and mounted using EuKitt.

### Quantification of Ki-61

Ki-67-positive cells were counted manually in the prosomere 2 region in E12.5 brain sections from control and *Tcf7l2^-/-^* animals from three different litters (3 mice, 4 sections each per genotype). Areas of each prosomere 2 section were measured in ImageJ and the number of Ki-67-positive cells by 1 mm^2^ was calculated. Two-tailed Student’s t-test was used to test for the difference between two groups (after confirming normal distribution of the data).

### DiI axon tracing

PFA-fixed brains were separated into hemispheres and small DiI crystals (Cat. no. D-3911, ThermoFisher Scientific) were placed in the exposed thalamic surface. Tissue was then incubated in 4% PFA at 37°C for 18-21 days. The hemispheres were then embedded in 5% low-melting-point agarose and cut into 100 μm thick coronal sections in a vibratome. The sections were counterstained with Hoechst, mounted onto glass slides, and secured under a coverslip with Vectashield Antifade Mounting Medium.

### Western blot analysis

Protein extracts were obtained from the thalami homogenized in ice-cold RIPA buffer. Protein concentrations were determined using Bio-Rad protein assay (Cat. no. 5000006, Bio-Rad Laboratories). Clarified protein homogenate (50 μg) was loaded on 10% SDS-polyacrylamide gels. Separated proteins were then transferred to Immun-Blot PVDF membranes (Cat. no. 1620177, Bio-Rad Laboratories), which were then blotted with anti-TCF7L2 (Cat. no. 2569, Cell Signalling), anti-β-actin (Cat. no. A3854, Sigma-Aldrich) and anti-GAPDH (Cat. no. SC-25772, SantaCruz) antibodies. The staining was visualized with peroxidase substrate for enhanced chemiluminescence (ECL) and 200 μM coumaric acid. Images were captured using Amersham Imager 600 RGB (General Electric).

### RNA isolation

*Tcf7l2*^-/-^ and WT littermate mice were collected on E18.5, and *Cck^Cre^:Tcf7l2^fl/fl^* and *Cck^Cre^:Tcf7l2^+/+^* mice on P60. Prosomere 2 regions (E18.5) or thalamic (P60) were dissected-out immediately and the RNA was extracted using QIAzol (Cat. no. 79306, Qiagen) and the RNeasyMini Kit (Cat. no. 74106, Qiagen).

### RNA-seq analysis

The quality of RNA was verified with Bioanalyzer (Agilent). RNA samples from three independent biological replicates for each genotype were sequenced on the same run of Illumina HiSeq2500. The reads were aligned to the mouse genome mm10 assembly from UCSC, using HISAT (Kim et al., 2015) and their counts were generated using HTSeq (Anders et al., 2015). Differential gene expression analysis was performed with DeSeq2 method (Love et al., 2014). Genes with log_2_(FC) > 0.4 and log_2_(FC) < −0.4 and FDR adjusted *p* value (*q* value) < 0.05 were considered to be the differentially expressed up- and downregulated genes.

### Functional enrichment analysis

Gene ontology enrichment was performed using an open access online tool GOrilla (http://cbl-gorilla.cs.technion.ac.il/). Two unranked lists of genes, one target and one universal (all detected transcripts) were used, assuming a hypergeometric distribution of the unranked genes. GO-Term enrichments were tested with Fisher’s exact test, and Bonferroni corrected *q* values <0.05 were considered significant.

### ChIP-Seq

C57BL mice were collected on P60 and sacrificed by cervical dislocation. Thalami were dissected, cut and fixed for 10 minutes in 1% formaldehyde (Cat. no. 114321734, Chempur), followed by 10 minutes of quenching with stop solution (Cat. no. 53040, Activ Motif). Chromatin was isolated from cellular nuclei according to (Cotney and Noonan, 2015), and then sonicated into 200-500 bp fragments with Covaris S220. Samples containing 10 µg of DNA were immunoprecipitated with 375 ng (10 µl) of TCF7L2 antibody (Cat. no. 2569, Cell Signaling). Incubation with protein G agarose beads and elution of precipitated chromatin was conducted according to ChIP-IT High Sensitivity kit protocol (Cat. no. 53040, Activ Motif). Cross-links were reversed and both eluates and input controls were treated with RNAse and Proteinase K, according to (Cotney and Noonan, 2015). DNA from 2 biological replicates was purified with Monarch PCR & DNA Cleanup Kit (Cat. no. T1030, NEB). Library preparation, sequencing and data analysis was conducted by NGS Core Facility of the Centre of New Technologies, University of Warsaw. Libraries were prepared with KAPA HyperPrep Kit and KAPA Dual-Indexed adapters (Cat. no. 7962363001, KK8722, Kapa Biosciences). Enrichment by 15 cycles of amplification was applied and final library was size-selected using Kapa Pure Beads to obtain average size of 350 bp. Libraries were tested for quality with High Sensitivity DNA kit (Cat. no. 5067-4626, Agilent) and then were sequenced on Illumina NovaSeq 6000 instrument in pair-end mode: 2×100 cycles. Raw reads were trimmed with the use of Trimmomatic software (Bolger et al., 2014) and then mapped to the reference genome, mm10 (UCSC) with the use of BWA (Li and Durbin, 2009). Duplicated reads were identified and removed with the use of Picard tool (Pentland et al., 1994). Both replicates for TCF7L2 and Input were merged into separate files, what resulted in 96% and 98% of properly paired reads, respectively. Peak calling was performed with the use of MACS2 (Zhang et al., 2008). Input was treated as a Control. Peaks were assigned to genes with annotatePeak function from ChIPseeker package using UCSC (University of California, Santa Cruz) annotation database. EdgeR package (Robinson et al., 2010) was used for statistic differential binding analysis between Input and TCF7L2 sample. To identify the most significant peaks we filtered the data for p(FDR) < 0.01, what resulted in 14 801 most significant peaks represented by 6 728 unique genes. At last we filtered the data for fold enrichment over the Input control ≥ 10, what resulted in 4625 peaks represented by 3791 unique genes. For de novo motif discovery the MEME-ChIP suite was used with DNA HOCOMOCO Mouse (v11 CORE) database (Kulakovskiy et al., 2013; Ma et al., 2014). Motif analysis was performed on filtered peaks, p(FDR) <0.01.

### In vitro slice electrophysiology

Brain slices (300 μm thick) from control and *Cck^Cre^:Tcf7l2^fl/fl^* mice (4 animals per genotype) of both sexes on P21-23 were prepared by an “along-row” protocol in which the anterior end of the brain was cut along a 45° plane toward the midline (Ying and Goldstein, 2005). Slices were cut, recovered and recorded at 24°C in regular artificial cerebrospinal fluid (ACSF) composed of: 119 mM NaCl, 2.5 mM KCl, 1.3 mM MgSO4, 2.5 mM CaCl2, 1 mM NaH2PO4, 26.2 mM NaHCO3, 11 mM glucose equilibrated with 95/5% O2/CO2. Somata of thalamocortical neurons in the ventrobasal complex were targeted for whole-cell patch-clamp recording with borosilicate glass electrodes (resistance 4-8 MΩ). The internal solution was composed of: 125 mM potassium gluconate, 2 mM KCl, 10 mM HEPES, 0.5 mM EGTA, 4 mM MgATP, and 0.3 mM NaGTP, at pH 7.25-7.35, 290 mOsm. Patch clamp recordings were collected with a Multiclamp 700B (Molecular Devices) amplifier and Digidata 1550A digitizer and pClamp10.6 (Molecular Devices). Recordings were sampled and filtered at 10 kHz. Analysis of action potentials was performer in Clampfit 10.6. Intensity to Voltage (I-V) plots were constructed from a series of current steps in 40 pA increments from −200 to 600 pA from a holding potential of −65 mV or at the resting membrane potential (around −57 mV). Two-tailed Mann-Whitney test was used to test for the difference in resting membrane voltage, membrane capacitance, numbers of action potentials and spiking frequency. Two-tailed Student’s t-test was used to test for the difference in series resistance (after confirming normal distribution of the data).

## Acknowledgements

We thank the Genomics Core Facility at the Centre of Genomic Regulation (CRG, Spain) and the NGS Core Facility at the Centre of New Technologies (NGSCF-CeNT, University of Warsaw) for providing next-generation sequencing services. Sequencing at the NGSCF-CeNT UW was done with the use of NovaSeq 6000 platform financed by the Polish Ministry of Science and Higher Education (decision no. 6817/IA/SP/2018 of 2018-04-10). We gratefully acknowledge M. Irimia from CRG, K. Misztal from the International Institute of Molecular and Cell Biology (Poland), K. Debski from the Nencki Institute of Experimental Biology (Poland), and D. Adamska, A. Mlodzińska and K. Goryca from NGSCF-CeNT for help with the RNA-seq analysis. We thank K. Brzozowska in our Laboratory for the initial management of the mice colony, and Kacper Posyniak for assistance with sample staining and acquisition. A part of genotyping was performed at the Laboratory of Animal Models facility at the Nencki Institute of Experimental Biology.

## Data availability

The RNA-seq raw FASTQ files are available at the EMBL-EBI data repository – ArrayExpress, under E-MTAB-8755 number. The ChIP-seq files are in the process of uploading at the NCBI Gene Expression Omnibus data repository. Their number will be provided as soon as the datasets are accepted.

---

**Table S1. RNA-seq analysis data from E18.5 WT and Tcf7l2^-/-^ thalami.** DeSeq2 list of all differentially expressed genes between E18.5 WT vs *Tcf7l2^-/-^*. List of all differentially expressed genes with a significant (p<0.05) log2 fold-change > 0.4 or < −0.4 between E18.5 WT vs *Tcf7l2^-/-^*.

**Table S2. Gene ontology enrichment analysis on the DEGs in thalami from E18.5 WT and Tcf7l2^-/-^ mice**. Reported GO terms (p≤0.001) divided into 3 ontologies: biological process, molecular function and cellular component.

**Table S3. RNA-seq analysis data from P60 WT and *Cck^Cre^:Tcf7l2^fl/fl^* thalami.** DeSeq2 list of all differentially expressed genes between E18.5 WT & *Tcf7l2^-/-^* and P60 WT & *Cck^Cre^:Tcf7l2^fl/fl^*. List of all differentially expressed genes between P60 WT & *Cck^Cre^:Tcf7l2^fl/fl^*. List of all differentially expressed genes with a significant (p<0.05) log2 fold-change > 0.4 or < −0.4 between P60 WT & *Cck^Cre^:Tcf7l2^fl/fl^*.

**Table S4. Gene ontology enrichment analysis on the DEGs in thalami from P60 WT and Cck^Cre^:Tcf7l2^fl/fl^ mice.** Reported GO terms (p≤0.001) divided into 3 ontologies: biological process, molecular function and cellular component.

**Table S5. TCF7L2 ChIP-Seq data in P60 thalami from WT mice.** 14 801 most significant peaks p(FDR) < 0.01 represented by 6 728 unique genes; 4 624 peaks with FC >10; Intersection of ChIP-Seq peaks with DEGs from RNA-Seq

**Table S6. Gene ontology enrichment analysis on the genes identified in TCFL2 ChIP from P60 thalami from WT mice.** Reported GO terms (p≤0.001) divided into 3 ontologies: biological process, molecular function and cellular component.

